# Spatial transcriptomic profiling reveals distinct signatures in acute versus chronic wounds through hypergraph modelling and transcriptomic entropy analysis

**DOI:** 10.1101/2025.02.21.639341

**Authors:** Megan C Sharps, Jeremy A. Herrera, Roxana Moscalu, Lily I Wright, Terence Garner, Syed Murtuza Baker, Adam Reid, Adam Stevens, Jason Wong

**Affiliations:** Tommy’s Maternal and Fetal Health Research Centre, Faculty of Biology, Medicine & Health, 5th Floor St. Mary’s Hospital, The University of Manchester, Manchester, UK; Wellcome Trust Manchester Cell-Matrix Centre, Faculty of Biology, Medicine and Health, The University of Manchester, Manchester, UK; Blond McIndoe Laboratories, Division of Cell Matrix Biology and Regenerative Medicine, School of Biological Sciences, Faculty of Biology, Medicine and Health, The University of Manchester, Manchester Academic Health Science Centre, Manchester, UK; Bioinformatics Core Facility, Faculty of Biology, Medicine and Health, The University of Manchester, UK; Department of Plastic Surgery & Burns, Wythenshawe Hospital, Manchester University NHS Foundation Trust, Manchester Academic Health Science Centre, Manchester, UK

## Abstract

**Introduction:** The financial burden of wounds to healthcare systems continues to increase, with little progression on advanced treatment options. Single cell RNA sequencing studies have predominantly focused on phenotypically different cell types and are generally descriptive in their analyses. Coordination within the transcriptome can be modelled using hypergraphs, which quantify higher order interactions (coordination between two or more genes) in transcriptomic data. Hypergraph modelling and analysis of transcriptomic entropy of wound samples will further elucidate differences between healing and non-healing wounds.

**Methods:** Spatial transcriptomic analysis (Visium Spatial Gene Expression) was performed on three acute wounds, three chronic wounds and two healthy, unwounded skin samples. Standard transcriptomic analyses were performed in Python using the packages Scanpy and Squidpy. Hypergraph modelling and transcriptomic entropy analyses were performed in R.

**Results:** Across the samples, twenty-nine Leiden clusters were defined, with keratinocytes, fibroblasts and adipocytes subclusters present. Fifteen marker genes for acute and chronic wounds were identified, with pathway analysis identifying immune responses and cellular homeostasis present in acute wounds, whilst in chronic wounds, pathways included immune responses and extracellular matrix structure. Spatial transcriptomics allowed for a descriptive analysis of the Leiden clusters within the samples, with each tissue sample being divided up into upper, middle and lower layer for this purpose. Hypergraph analyses of the whole transcriptome from the pathology groups and the independent transcriptomic Leiden clustering, revealed differences in underlying transcriptomic coordination. The top 1000 highly connected genes from each pathology group were analysed by Over Representation Analysis, with integrated stress response signalling, maintenance of cell number and myeloid and mononuclear cell differentiation pathways present in acute samples (all FDR <0.05) compared to regulation of vasculature development, mononuclear cell differentiation and epithelial cell proliferation (all FDR <0.05) in the chronic samples. Analysis of transcriptomic entropy identified further differences between acute and chronic wounds, with decreasing entropy from control to acute to chronic samples (all p-adjusted <0.001). When comparing Leiden clusters that were present in at least two sample groups, there was a statistically significant difference in entropy, although Leiden clusters varied in high or low entropy between acute and chronic wound samples (all p< 2.2 x 10^−16^). By mapping high and low entropy values to spatial images of each sample, lower transcriptomic entropy within the wound bed was identified compared to the rest of the tissue sample.

**Conclusions:** Spatial transcriptomic analysis allowed for distinct spatial patterning to be observed which delineated acute and chronic wounds. Analysis of higher order interactions within the wound sample transcriptome identified key pathways that were not identified using traditional analyses. Transcriptomic entropy analysis further highlighted the differences between the wound samples. This work has implications for the future development of biomarkers of chronic wounds.

## Introduction

The burden of wounds on society costs the NHS £8.3 billion a year in the UK as of 2017/2018, with costs likely to still be rising, and increased costs of over $20 Billion a year in the USA [1, 2]. Globally, wounds are a growing multibillion dollar industry that is set to rise by $30+ Billion by 2029 [3]. In the UK, the cost of treating a wound to heal can be up to £4684, and can increase to £7886 for a chronic, non-healing wound [1]. Whilst acute wounds have been carefully characterised in both animal and human models, we still fundamentally do not know what the differences between acute and chronic non-healing wounds are other than through clinical aetiology mechanisms derived from accurate history taking and imprecise investigations such as doppler wave forms, microbiological swabs and basic histology to exclude specific pathologies such as skin cancer [4]. The other limitation of studying chronic wounds is the lack of any *in vitro* and *in vivo* models with specific human relevance. *In vitro* models such as 3D cell cultures [5] are contributing to the reduction on the dependence on animal models, but cannot yet mimic the critical elements of cellular infiltration. Models used to mimic wound healing including murine models of oxidative stress [6] and aged diabetic mice [7] capture the importance of systemic cellular contributions but simply cannot mimic the complex heterogeneity of conditions that allow for an informative basis to treat the disease. As such, attention has been focused on deriving treatment efficacy directly on patients. This has led to the massive proliferation of wound care products that lack mechanistic evidence of efficacy but are continued in use as there are no alternatives to this massive healthcare burden [8].

The exact categorisation of the stages of wound healing following an injury to the skin have been under debate. The original paradigm was composed of five main stages; haemostasis, inflammation, growth, re-epithelialisation and finally, tissue remodelling [9]. This process has been well defined in several species, resulting in vasoconstriction and the formation of a platelet plug during haemostasis immediately following injury, the recruitment and actions of immune cells during the inflammatory stage, neovascularisation and proliferation of keratinocytes in the growth and re-epithelialisation stages respectively and finally continued tissue remodelling following closure of the wound. However, whilst there are hypotheses as to why impairment of wound healing occurs resulting in chronic wounds, such as prolonged inflammation [10] and prolonged protease activity [11], the exact mechanistic pathways which result in a wound becoming chronic is yet to be elucidated. Following spatial analyses of wounds, a new three stage paradigm has been proposed [12]. The first proposed stage is early inflammation, with the recruitment of immune cells. This is followed by proliferation of both macrophages and keratinocytes, and recruitment of fibroblasts and migration of keratinocytes leading to reepithelization. The final stage is the activation of fibroblasts leading to fibrosis. This simplified paradigm may influence how wounds are treated in clinical practice.

The main difficulty with investigating complex wounds is obtaining appropriate wound samples for analysis, with the heterogeneity between wound types, such as area, depth and time since injury of the wound, alongside patient co-morbidities, such as diabetes, vascular disease and age, all contributing to the failure of the wound healing process [9, 13]. However, despite the aetiologies being different, we hypothesise that chronic wounds will have a number of similarities that preclude them from healing due to a perturbed immune and matrix biology. This theory can only really be tested by the relevant chronic wound model or from real patient data. To date, investigations of the biology of complex wounds have been focused on traditional histochemistry, with more recent studies using single cell RNA sequencing (scRNA) and FISH technologies [14–16]. Whilst scRNA data has provided a number of important insights into wound healing biology in particular cellular heterogeneity in the epithelium [14] and fibroblast phenotypical evolution [16] in resolving wounds, or a single chronic wound type [16], the lack of spatial data is also hampering the ability to deconvolve the biology of complex wounds as scRNA typically doesn’t allow for the contextual anatomy of the deeper layers of tissue to be examined, or for the spatial location of the cell types to be determined.

To date, largely descriptive publications have allowed for qualitive analysis describing differential gene expression over a map of cell types. However, a more quantitative comparison of coordination in the transcriptome is possible using hypergraphs. Traditional networks model pairwise interactions either between or within groups [17]. A traditional network approach can miss features of the transcriptome such as coordination that are important for function in a complex system. Hypergraphs allow for the interaction between two or more nodes to be analysed, known as higher-order interactions, and therefore provides a quantitative way to define a function model of the transcriptome in acute and chronic wounds [18]. Transcriptomic entropy can be calculated over the hypergraph, thereby providing information as to the level of order or disorder with a system [19].

The current wound healing knowledge is growing exponentially with techniques such as scRNA and spatial analysis in a variety of models. To date, there is a plethora of high-quality data from wound samples, both human and animal models [12, 20, 21], however, much of the analysis has been focussed on identifying cell types of interest and phenotypic changes that occur during the wound healing process. The importance of understanding the context of these findings in relation to real human chronic pathologies historically has relies upon fresh samples. The ability to analyse FFPE samples by Visium Spatial Gene Expression allows for simple logistical barriers to be overcome.

To address the dearth in understanding we have initiated an analysis of normal skin, acute and chronic wounds from a range of patient tissues accessed through our biobank. This has provided us with tissue sources that have detailed clinical information, and standardised harvesting techniques allowing for detailed spatial transcriptomic characterisation using the Visium Spatial Gene Expression platform. We hypothesise that by analysing higher order interactions and entropy within spatial transcriptomic data will help elucidate the differences between acute and chronic wounds. These analyses will provide a framework on which future biomarkers of chronic wounds can be identified.

## Methods

### Sample collection

Biopsy samples were collected at the time of surgery, all patients consented before being taken to theatre and samples were collected by a biobank nurse. Standardised surgical biopsies were taken based on a 1cm^3^ excision at the wound edge, or non-injured site for control tissues. Samples were washed in 0.9% saline before being fixed in 10% buffered formalin for 24-48 hours at room temperature and stored in 70% ethanol until processed into formalin-fixed paraffin-embedded (FFPE) blocks. Samples were identified as being acute, chronic, or controls. All samples were collected following informed consent and all procedures were approved by the National Health Research Authority through the ComplexWounds@Manchester Biobank (NRES 18/NW/0847).

Confirmation of the skin and wound characteristics were made with a histopathologist, including outlining the epidermis, papillary dermis, reticular dermis, hypodermis and wound bed.

### Spatial Transcriptomics

#### Tissue RNA Quality Assessment

To determine appropriateness of the samples for Visium Spatial Gene Expression for FFPE (V1, 10X Genomics) analysis, the quality of the RNA was determined following extraction using RNeasy FFPE Kit for RNA isolation (Qiagen, 73504; manufacturers recommendation) from 2-4 sections of tissue per sample cut at 10 microns and then eluted in 15 µL of water. We followed manufacturers recommendation to assess RNA quality by mixing 5 µL of RNA sample with 1 µL of High Sensitivity RNA ScreenTape Sample Buffer (Agilent; 5067-5580) and incubated at 72°C for 3 minutes. The sample was then loaded onto a High Sensitivity RNA ScreenTape (Agilent; 5067-5592) and analyzed on a 2200 Tapestation system (Agilent). We selected ‘Region Settings’ and entered the lower limit of 200 nucleotides and upper limit of 10,000 nucleotides to calculate a DV200 score. An acceptable score of 50% or higher was used to ensure that the sample was fit for Visium Spatial Gene Expression.

#### Visium Spatial Gene Expression

Using a ruler and razor blade, a 6 mm^2^ region of interest was carved out of the tissue block and re-embedded into a new FFPE block. The wax of the block was further trimmed so that a 2 mm border of wax surrounds the tissue to ensure proper sectioning and mounting onto the fiducial frame of the slide. Following 10X Genomics recommendations, the microtome blades were wiped with 100% ethanol and a 5 µM thick tissue section was placed into a 42°C clean water bath. Using a paintbrush, the section was oriented onto the fiducial frame. The slide was then dried in a 42°C incubator for 3 hours and stored in a desiccator at room temperature overnight. The following day, deparaffinization, haematoxylin and eosin staining was performed, followed by the addition of a coverslip, and imaging (Olympus BX63) following 10X Genomics recommended reagents and protocol (protocol CG000408). The coverslip was removed, and the tissue section was de-crosslinked overnight using a BioRad model thermocycler. The next day, the exact 10X Genomics protocol was followed for probe hybridization, ligation, RNA digestion and probe release and extension (protocol CG00407, RevC). For library construction, cDNA concentration was analysed by using a Biorad CFX96 to calculate a Cq value per specimen and followed 10X Genomics recommended reagents and protocol to create a library. Samples were cleaned up using SPRIselect and eluted in a volume of 25 µL and stored at −20C. Images were uploaded into Loupe Browser (V8, 10X Genomics) to align the image and calculate the coverage of tissue over the spots. The tissue coverage and cDNA libraries were given to the University of Manchester Genomics Core Facility for sequencing on an Illumina Next Seq500.

### Computational Analyses

#### Data processing

Fastq files for each sample were processed using the standard Space Ranger (V3.0.0) analysis pipeline using JSON alignment files generated from use of Loupe Browser (V8). Data were aligned to the GRCh38 reference genome, with datafiles for downstream analysis generated.

Analysis was performed in Python using standard packages such as Scanpy [22] and Squidpy [23]. Raw data were integrated into one Anndata file for downstream analysis. Cells with less than 100 genes expressed were removed alongside genes that were expressed in fewer than 3 cell spots. This was to ensure biological differences were maintained, whilst also ensuring quality control of the data. Data were then library normalised, log transformed and the PCA calculated using standard functions in Scanpy. Batch correction was performed using Harmony. Dimensionality reduction of the integrated dataset was performed and visualised using Uniform Manifold Approximation and Projection (UMAP). Clustering was performed using the Leiden function in Scanpy, with the resolution set to 1.7, with the tissue structure taken into consideration. Differential gene expression was performed using the rank_genes_group function in Scanpy, with a Wilcoxon rank-sum test. Statistical significance was set at p<0.05. Resulting differentially expressed genes were assessed using The Human Protein Atlas [24] and EnrichR [25] to determine cell types present within each cluster. Data were compared to both adipose and skin samples depending on the spatial location of the clusters.

#### Hypergraph and transcriptomic entropy analyses

Hypergraph and transcriptomic entropy analyses were conducted in R on data exported from Python. Two forms of hypergraph analysis were performed.

Hypergraphs are generalisations of graphs (networks) wherein an edge can connect an arbitrarily large number of nodes. A hypergraph was generated by correlating the genes of the transcriptome against themselves. The resulting matrix was binarized around the standard deviation of the correlation r values. In doing this, the incidence matrix of a hypergraph (*M*) is generated; the reduced adjacency matrix of the hypergraph (*MM^T^*) can be derived by multiplying this matrix by its transpose (*M^T^*). The row sums of the reduced adjacency matrix represent the connectivity of nodes within the network, whilst capturing the number of other nodes shared by those connected edges – a higher-order feature which distinguishes hypergraphs from pairwise networks. This was calculated for genes in each pathology group (acute, chronic or control) independently. Groups were ranked, averaging in the case of tied rank, and a normalised ranking was calculated by dividing the row sum by the total number of edges. The list of genes and their ranking were taken forward for Gene Set Enrichment Analysis.

A second, iterated, variation of the hypergraph analysis was performed to calculate transcriptomic entropy, using the entropy package in R. This was performed in two ways: 1) generating a single hypergraph per pathology group and 2) a single hypergraph per cluster per pathology group. A threshold was set so that a cluster had to be present in >90 cell spots to be included in the data of the pathology group, to eliminate clusters with limited data. To enable direct comparisons between clusters and the different pathology groups, the correlation matrix underlying the hypergraph was generated from a randomly selected set of 100 genes, correlated against an additional, random subset of 1000 genes. The hypergraph was iterated 10,000 times to ensure robustness of the data, with the entropy of the reduced adjacency matrix of the hypergraph calculated for every iteration. Data were plotted as box plots to compare pathology groups, violin plots containing boxplots to compare clusters within each pathology group, as well as data grouped into “high” (median entropy ≥9) or “low” (median <9) entropy of each cluster-level entropy and plotted on an image of the samples to investigate spatial patterns.

#### Mapping transcription factors

To determine if any of the top 1000 row sum distribution ranked genes were transcription factors, a list of 881 known transcription factors was downloaded from the Factorbook web-based repository [26]. The top 1000 genes per pathology group were compared to the list of transcription factors and the results plotted in a Venn diagram and gene ontology performed. The transcription factors unique to each pathology group were combined into a dictionary in Python and plotted as a spatial heatmap.

#### Pathway enrichment analysis

Gene Set Enrichment Analysis (GSEA) was conducted using the compareCluster function in R, using gseGO. Genes were compared to the OrgDb for Homo sapiens, significance set to p<0.05 and pAdjustMethod set to Benjamini Hochberg. Firstly, all genes per pathology group were ranked using the row sum distribution ranking. Secondly, the top 1000 highly ranked genes per pathology group were ranked based on the row sum distribution ranking.

Gene ontology was performed on the list of transcription factors that mapped to the hypergraph genes for each pathology group using WEB-based GEne SeT AnaLysis Toolkit [27]. Over Representation Analysis was performed, using a Fisher’s exact test, using the genome-protein coding reference set.

#### Statistical analysis

Statistical analyses were performed on the transcriptomic entropy per pathology group, using a Wilcoxon with Bonferroni correction, with significance set at p<0.05. To calculate the statistical difference in the transcriptomic entropy distribution per cluster between pathology groups, a Wilcoxon was performed, with significance set at p<0.05

## Results

The demographics of the patients and the description of the wound samples can be found in Supplementary Table S1. The final spatial transcriptomic analysis was performed on three acute wounds, three chronic wounds and two control skin samples. Sixty two percent of samples were from female patients with 80% of the patients of White British ethnicity. The average age of acute wounds was 8 days, compared to 22 months for the chronic wounds.

The acute samples were harvested from several traumatic wounds where sampling was conducted between the time of injury and timing of reconstruction, typically within 10 days. The chronic wounds were defined as those where the wounds had failed to heal after 6 weeks [28, 29] and were from defined conditions related to failure of wounds to progress (pressure sores, diabetic foot ulcer, post traumatic wound). All the acute samples healed 6 months post-sampling, whilst none of the chronic samples had healed at 6 months post-sampling. Wounded samples were compared to healthy, unwounded skin.

### Microscopic analysis and histopathological annotation

The annotation of the defined tissue layers of each sample highlighted distinct differences between each group. Annotated images of each sample can be found in Supplementary Figure 1. Control skin had well defined anatomical structures relating to the epithelium, papillary dermis, reticular dermis and hypodermis adipose tissue readily perceivable. All samples in the acute wound group had an identifiable wound bed, epithelium, both papillary and reticular dermis layers, and adipose tissue. The wound edge was well defined and distinct anatomical structures were still identifiable. Areas of angiogenesis indicated by vascular channels filled with red blood cells, and cellular infiltration could be seen in the fat lobules. In contrast, the chronic wounds all had identifiable wound beds, and epithelium, but the papillary and reticular dermis were hard to discern, as the matrix was far denser and infiltrated with cells. Very little hypodermal fat could be identified in these samples.

### Spatial Transcriptomic analysis

After following standardised batch correction and quality control, 25,780 cells spots were available for analysis across the 8 samples. Twenty-nine cell spot clusters were identified, with cell types included in the clusters determined by gene expression and location of the cell spots within the tissue (Figure 1A). The proportion of Leiden clusters varied between the sample groups (Figure 1B&D). A correlation matrix between the whole transcriptome of all the samples identified that acute and control samples clustered together separately from chronic samples, likely due to the similarity in the tissue structure beneath the acute wound bed and control samples (Figure 1C).

**Figure 1:**
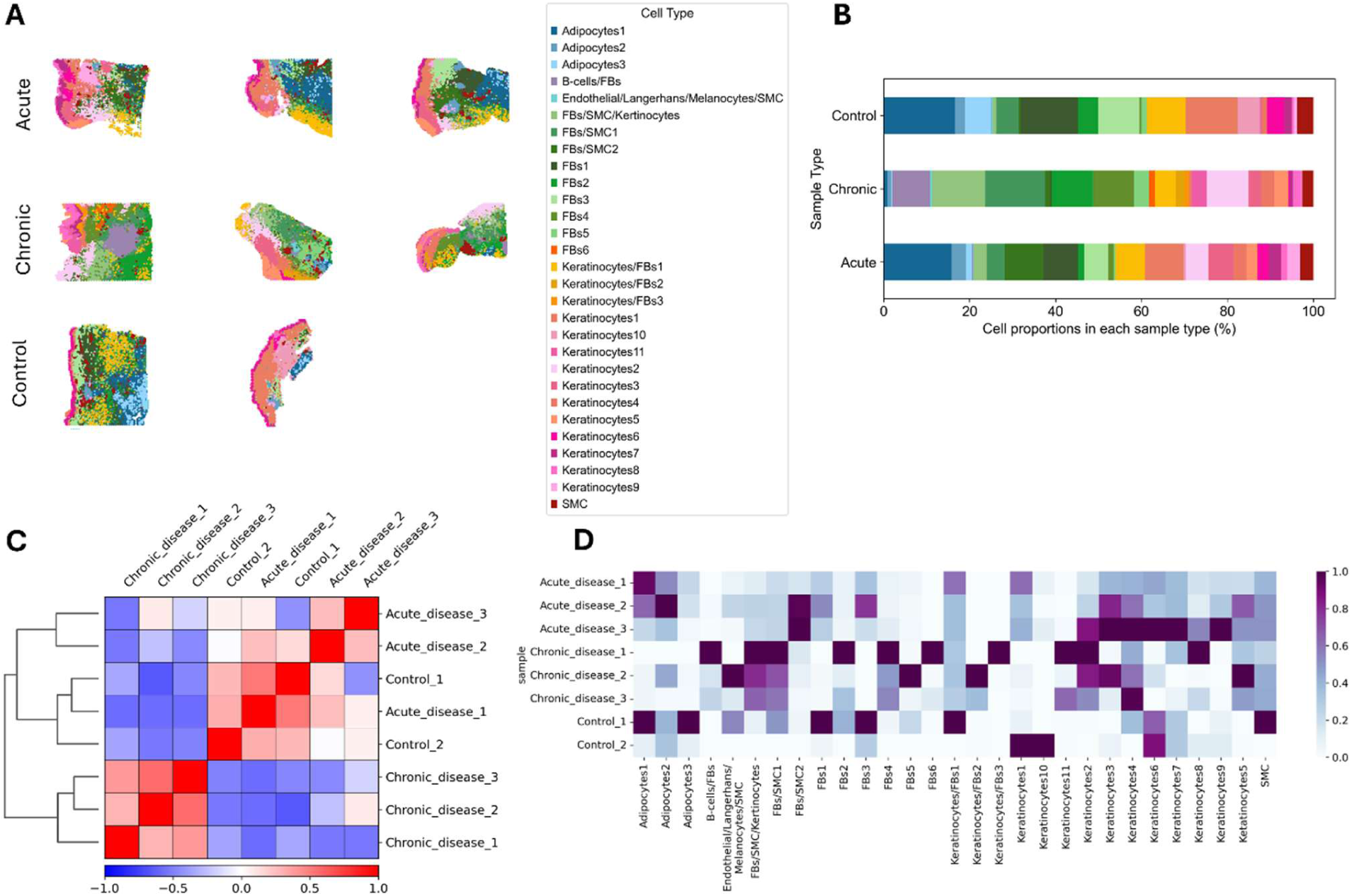
Visium Spatial Gene Expression analysis of Acute and Chronically (n=3/group) wounded skin and Control (n=2) skin samples. **A)** Spatial location of Leiden clusters on acute, chronic and control samples. **B)** Proportion of Leiden clusters in each pathology group. **C)** Correlation matrix of the similarities between each sample. Red indicates increased similarity, blue indicates less similarity. **D)** Heatmap of the expression of Leiden clusters per sample. FBs: Fibroblasts, SMC: Smooth Muscle Cells

Differential gene expression was conducted using the rank_gene_group function in Scanpy. Due to the size of the barcoded spots and therefore the resolution of the sequencing, Visium V1 cannot determine individual cell types. Therefore, to determine the main cell types within each cluster, the top differentially expressed genes were compared to data from EnrichR [25] and The Human Protein Atlas [24], alongside the spatial location of each cluster within the tissue samples. There were 11 keratinocyte only containing clusters, and an additional four mixed clusters containing keratinocytes. Six clusters only contained fibroblasts, with an additional seven mixed cell type clusters containing fibroblasts. Three clusters contained adipocytes and two contained immune cells. Mixed clusters in general contained fibroblasts, endothelial cells, smooth muscle cells, keratinocytes and immune cells (Figure 1A-D). A dot plot with the top 5 differentially expressed genes per cluster can be found in Supplementary Figure 2.

To identify marker genes for each pathology group, the top 15 differentially expressed gene per group with a significant p-value and a minimum positive fold change of 1 were investigated (Figure 2A). In all groups, the marker genes were identified as being present within the dermis, rather than at the wound bed in the chronic and acute groups (Figure 2B). Three of the chronic sample marker genes played a role in immune response (*IGKC, IGHG* and *KRT16*), whilst the remainder were linked to extracellular matrix (ECM) structure (including *COL1A1, FN1, BGN* and *POSTN*) (Figure 2A). In contrast, in acute wounds, three different genes were linked to the immune response (*C1QC, CEPBD* and *CFD*) whilst the remainder of the genes were linked to processes such as cell cycle progression (*MYC*), cellular response to stress (DUSP1) and cell mobility and survival (*HSPA5* and *MARCK5*). In control samples, the marker genes were linked to general homeostasis pathways (*KRT2, CACL12, FOS, GLUL* and *ADIRF*).

**Figure 2:**
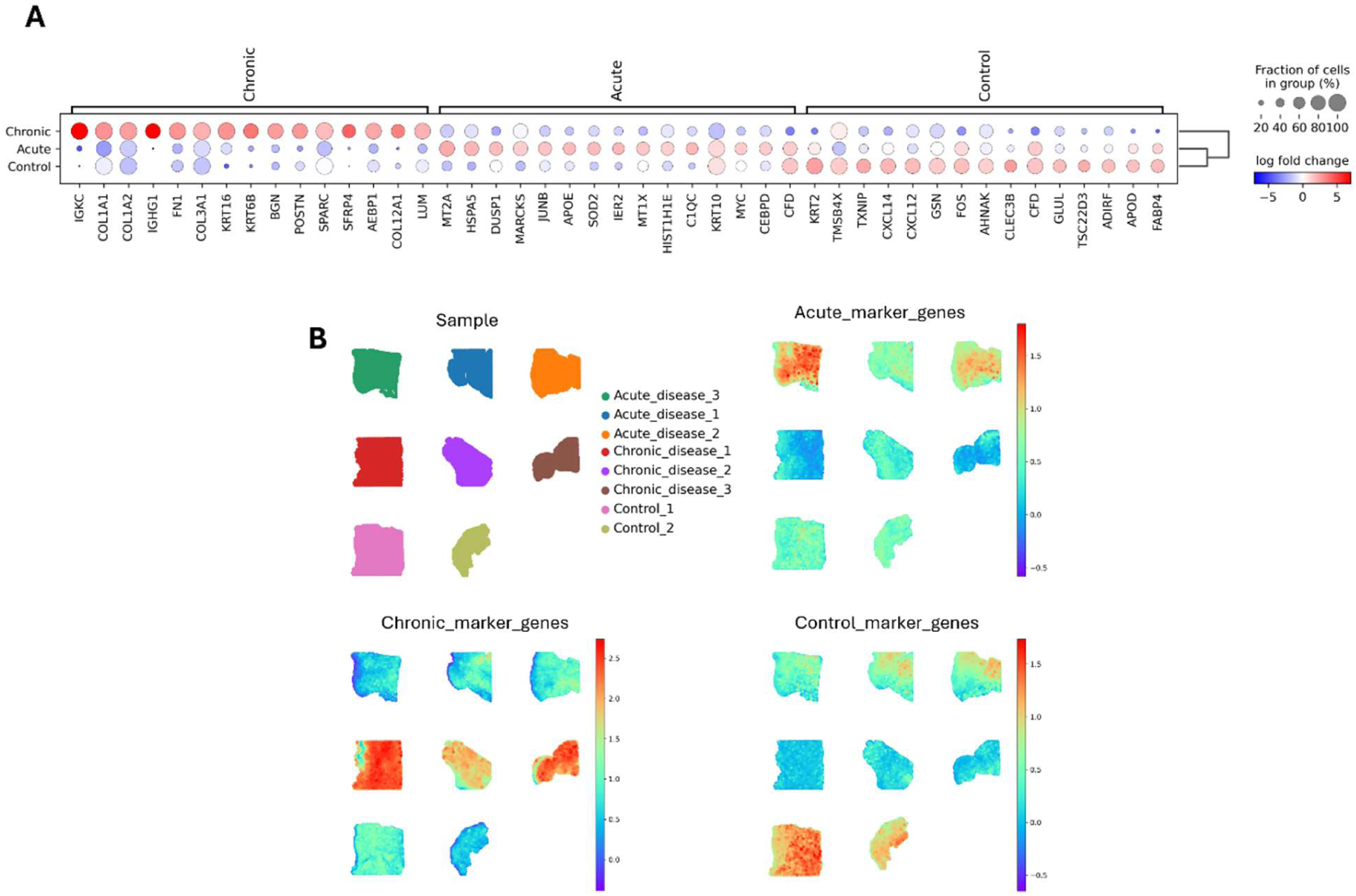
Transcriptomic markers of pathology groups. **A)** Dot plot of the top 15 genes with a positive log fold change >1 that differentiates Acute and Chronic (n=3/group) wounded skin samples and Control (n=2) skin samples. Genes determined by using the rank_gene_group function in Scanpy, and with a p value threshold of <0.05. **B)** Spatial location of transcriptomic markers of pathology groups. Top 15 genes were combined into a list before being plotted using the pl.spatial function in Scanpy.

### Comparison of cluster spatial locations between sample groups

Next, we identified the spatial location of samples (Figure 1A) and how they compared between pathology groups. This allowed us to take into consideration the defined layers of tissue from the H&E images (Supplementary Figure 1). We have defined the layers of tissue as upper, middle and lower for the purposes of the tissue description but have also presented quantitative information on the clusters present in each pathology group (expressed as cell spots per pathology group).

#### Control, unwounded samples

The quantitatively prevalent normal skin clusters include the following cell spot numbers: Keratinocytes1 (700), Keratinocytes6 (230), Keratinocytes10 (297), Fibroblasts1 (771), Fibroblasts3 (559), Adipocytes1 (959) and Adipocytes3 (351).

In control unwounded skin, there were clusters that relate to the spatial anatomy, which are generalist to the various layers of the skin. Across the levels of unwounded skin, we see:

##### Upper layer

The clusters in the upper layer relate to the keratinocyte clusters and were highlighted by clusters Keratinocytes1, Keratinocytes6 and Keratinocytes7. Fibroblasts3 relates to a subnormal fibroblast population. The Keratinocytes4 cluster was seen in one of the control samples but not the other. The clusters of Keratinocytes7 and Keratinocytes8 are very distinct for the epidermis, Keratinocytes4 was more expressed in the basal epidermis, whilst Fibroblasts3 and Keratinocytes10 were more subepithelial.

##### Middle layer

The clusters observed in the middle layer of the tissue all contain fibroblasts. Clusters Fibroblast/Smooth Muscle Cell1 and Keratinocytes/Fibroblasts1 were determined to be related to general dermal activity seen in skin. In the control samples, this cluster was found throughout the tissue, and there was generalised aggregation in the dermis of chronic wounds. In contrast, in the acute wounds, it was located at the wound edge. The cluster of Fibroblasts1 is also dominant across the dermis. There were similar expression patterns in the clusters Fibroblasts/Smooth Muscle Cells/Keratinocytes and Fibroblasts4, which were scantily seen in normal skin. Another sparsely observed cluster in control samples is the Fibroblasts/Smooth Muscle Cell2 cluster. Cluster Fibroblast2 is a deeper subset of fibroblasts, which probably relates to the reticular layer and hypodermis. Fibroblast5 has speckled expression across the dermis. Relating to the vascular patterning in the dermis is the Smooth Muscle Cell cluster whilst the Adipocytes2 cluster is found in the middle of the sample as opposed to the hypodermal region.

##### Lower layer

Below the dermis in the normal unwounded skin is cluster Adipocytes1 which relates to fat microvacuoles which are evident in the deeper layers of control skin. The cluster Adipocytes3 is a subtype of fat compartments in normal skin and is also present in acute wound samples. However, this cluster is sparsely found in chronic wounds. The cluster Endothelial cells/ Langerhans/Melanocytes are sparsely expressed in the lower dermis.

#### Acute wound samples

The quantitatively prevalent clusters seen in acute wounds include the following cell spot numbers: Keratinocytes1 (887), Keratinocytes3 (587), Keratinocytes4 (310), Keratinocytes5 (248), Keratinocytes6 (263), Keratinocytes7 (295), Keratinocytes9 (298), Fibroblasts1 (807), Fibroblasts3 (572), Adipocytes1 (1584), Adipocytes2 (345), Keratinocytes/Fibroblasts1 (693) and Fibroblasts/Smooth Muscle Cells2 (903).

The highly specific clusters for acute samples include Keratinocytes1, Keratinocytes6, and Keratinocytes7 for keratinocyte layers, Fibroblasts1 and Fibroblasts3 for fibroblast-related clusters and Adipocytes1 and Adipocytes3 for adipocyte clusters. The Fibroblasts/Smooth Muscle Cells2 cluster is also specific for acute wound samples. Across the levels of acute wounds, we see:

##### Upper layer

At the leading edge of the wound is Keratinocytes5 followed behind by a large focus of Keratinocytes3. Cluster Keratinocytes6 sits on the very upper aspect of the skin as a thin single layer of epidermis. A slightly thicker layer of cluster Keratinocytes7 is also within this region. The cluster Keratinocytes1 demonstrates a confluent sub-epidermal or basal epidermal layer of activity. Finally, close and slightly deeper to this cluster are Keratinocytes9 and Keratinocytes10 and behind and deep to this is cluster is Keratinocytes4.

##### Middle layer

The cluster Keratinocytes2 sees a focus of cells underneath the leading edge of the epidermis in acute wounds. Fibroblasts3 are seen in the mid-dermis away from the wound edge, whilst Fibroblasts4 is seen as scanty expression in the middle of the sample. Adipocytes2, a discrete adipocyte cluster, sits in the mid-dermis region. Cluster Fibroblasts/Smooth Muscle Cells2 shows dense expression of the middle of the samples.

##### Lower layer

The Adipocyte1 cluster can be seen in acute wounds, along with Fibroblasts/Smooth Muscle Cells1 in the lower adjacent wound area. When looking spatially at the samples, there are clear layers of clusters as you move from the wound deeper into the tissues. The order of the clusters starts with Keratinocytes/Fibroblasts to Fibroblasts/Smooth Muscle Cells1 and Fibroblasts/Smooth Muscle Cells/Keratinocytes and Fibroblasts2, into Adipocytes1, Fibroblasts1, Fibroblasts5, and Adipocytes3 clusters.

#### Chronic wound samples

The quantitatively prevalent clusters seen in chronic wounds include the following cell spot numbers: Keratinocytes2 (968), Keratinocytes4 (310), Keratinocytes5 (325), Keratinocytes8 (206), Keratinocytes11 (343), Fibroblasts2 (959), Fibroblasts4 (946), Fibroblasts5 (347), Fibroblasts6 (142), Fibroblasts/Smooth Muscle Cells1 (1386), Fibroblasts/Smooth Muscle Cells/Keratinocytes (1235), B-cells/Fibroblasts (897), Keratinocytes/Fibroblasts2 (196), Keratinocytes/Fibroblasts3 (103), Endothelial cells/ Langerhans/ Melanocytes/ Smooth Muscle Cells (48).

There are five clusters that are highly specific to the chronic wound samples. There are the Keratinocytes11, Fibroblasts10, B-cell/Fibroblasts, Keratinocytes/Fibroblasts2 and Keratinocytes/Fibroblasts3 clusters. The spatial anatomy of the chronic wound shows:

##### Upper layer

There is a marked decrease in expression of the following keratinocyte clusters: Keratinocytes1, Keratinocytes3, Keratinocytes6, Keratinocytes7 and Keratinocytes10, but an increase in Keratinocytes11, Keratinocytes/Fibroblasts2 and Keratinocytes/Fibroblasts3. The Keratinocytes2 cluster of keratinocytes are seen expressed in the wound bed.

##### Middle layer

Fibroblasts/Smooth Muscle Cells1 has dense expression throughout the mid to lower skin level and the B-cells /Fibroblasts cluster can be seen as a condensation of expression in the middle of the samples. This is a different expression that in acute samples (897 cell spots vs 19 cell spots in chronic wound vs acute wound samples). Interestingly, there is an absence of expression of both Fibroblasts3 and Adipocytes2 clusters. The Fibroblasts4 cluster is more prominent in chronic wounds in the upper wound dermis and Fibroblasts6 is found in chronic wound samples but not in acute wound samples. The Fibroblasts/Smooth Muscle Cells/Keratinocytes mixed cluster is also seen in greater number in the chronic wound samples compared to the acute wound samples (1235 vs 302 cell spots) and represents the wound bed.

##### Lower layer

Adipocytes1 has very low expression in chronic samples compared to controls and acute samples (92 vs 959 vs 1584 cell spots, respectively). The cluster Fibroblasts1 are much less in chronic wounds, whilst in contrast, Fibroblasts2, Fibroblasts5, and Fibroblasts6 clusters are seen in the lower dermis and expressed greater in chronic wounds. Additionally, Adipocytes3 expression is reduced in chronic wounds.

An interesting observation was that of the Smooth Muscle Cell cluster that was seen across all pathology groups. Despite being present in relatively similar numbers of cell spots (307, 256 and 217 cell spots in acute, chronic and control samples, respectively), the degree of entropy in these clusters across pathology differed significantly (Figure 5E).

### Presence of known marker genes of pro- and anti-inflammatory states, immune cells, angiogenesis, proliferation and apoptosis

Known markers of cell types and reported mechanisms of wound healing were collated (Supplemental Table S2) and compared for differences in spatial patterning. These included pro- and anti-inflammatory markers, proliferation markers, angiogenic markers, apoptotic markers, Langerhans cells, macrophage phenotypes, dendritic cells, T cells and B cells were compared to our samples (Figure 3A-P).

**Figure 3:**
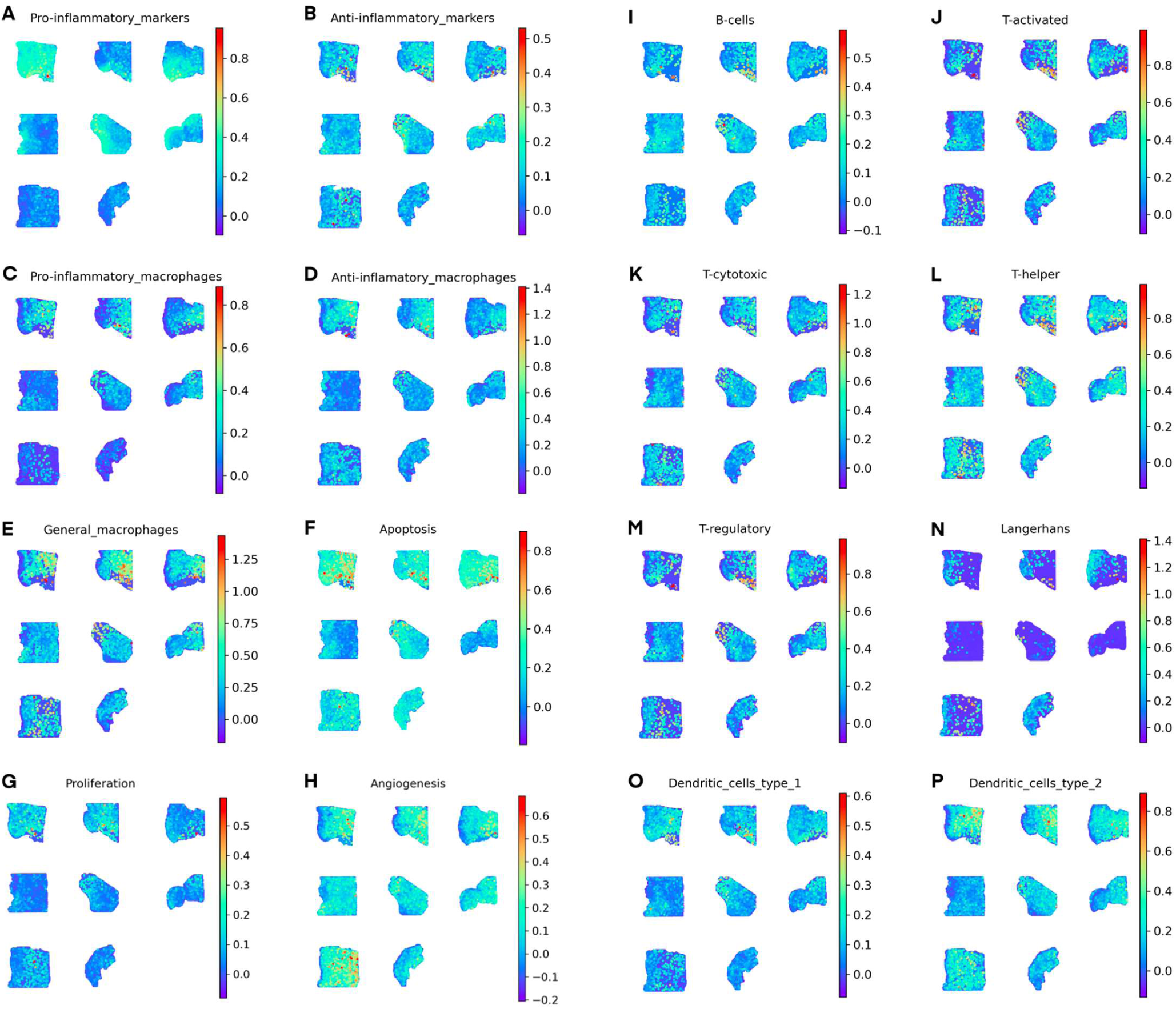
Spatial location of known markers of pro- and anti-inflammatory markers and immune cells. **A)** Pro-inflammatory markers, **B)** Anti-inflammatory markers, **C)** Pro-inflammatory macrophage markers, **D)** Anti-inflammatory macrophage markers, **E)** General macrophage markers, **F)** Apoptosis markers, **G)** Proliferation markers, **H)** Angiogenesis markers, **I)** B-cell markers, **J)** T-activated cell markers, **K)** T-cytotoxic cell markers, **L)** T-helper cell markers, **M)** T-regulatory cells markers, **N)** Langerhans cell markers, **O)** Dendritic cell type 1 markers, **P)** Dendritic cell type 2 markers. Markers used in the generation of the plots can be found in Supplementary Table S2 with genes saved as a lists before plotting using the pl.spatial function in Scanpy.

There were spatially distinct patterns in the acute and chronically wounded skin. In both the acute and chronic samples, there was an increase in the pro-inflammatory markers compared to controls. Two of the three acute wounds had pro-inflammatory macrophages present in the wound bed, compared to all three chronic samples. In contrast, in acute wounds there was a general absence of anti-inflammatory markers in the wound bed, apart from the occasional highly expressing cell spot, whilst in the chronic samples, anti-inflammatory markers were present more uniformly. There were sporadic cell spots containing general macrophage markers in the areas of the acute wound beds alongside T regulatory cells. There was also an increase in both apoptosis and proliferation compared to both chronic wounds and control samples. In chronic wound samples, there was a reduction in both anti-inflammatory macrophages and Langerhans cells compared to acute wound and control samples. However, where there was presence of anti-inflammatory macrophages, they were generally near the wound bed. Across all three groups, there was uniform distribution of B-cells, T-cytotoxic cells and T-helper cells, alongside dendritic cells and angiogenesis markers, with some areas with higher expression within each sample.

### Hypergraph and transcriptomic entropy analysis

Hypergraph and transcriptomic entropy analysis was performed to investigate the coordination within and organisation of the transcriptome in acute and chronic wounds and healthy control skin.

#### Hypergraph analyses

Hypergraph analyses allow for the coordination within the transcriptome to be investigated and identified key functional pathways in pathology groups. When considering the network structure of all genes in the transcriptome in a GSEA analysis, pathways involved in keratinisation, RNA splicing and cell-substrate adhesion were present across both pathologies and the control samples (Figure 4A). However, when the top 1000 highly connected genes were investigated, some pathways remained common between the three sample groups, whilst differences between the acute and chronic samples also became apparent (Figure 4B). Pathways identified in the GSEA of the top 1000 ranked genes included peptide cross-linking, intermediate filament cytoskeleton organisation and intermediate filament-based processes were present across both pathologies and the control samples. There were also common pathways between acute wounds and control samples, including cell adhesion, and cellular developmental process, and between chronic wounds and control samples, such as keratinocyte differentiation, epidermal cell differentiation and epidermis development. The overlap between the top 1000 highly connected genes in the group hypergraphs were compared. Of the 1000 highly connected genes, 378 were shared between the three sample groups. A total of 275 genes were unique to acute samples, 357 were unique to the chronic samples and 334 were unique to the control group (Figure 4C).

**Figure 4:**
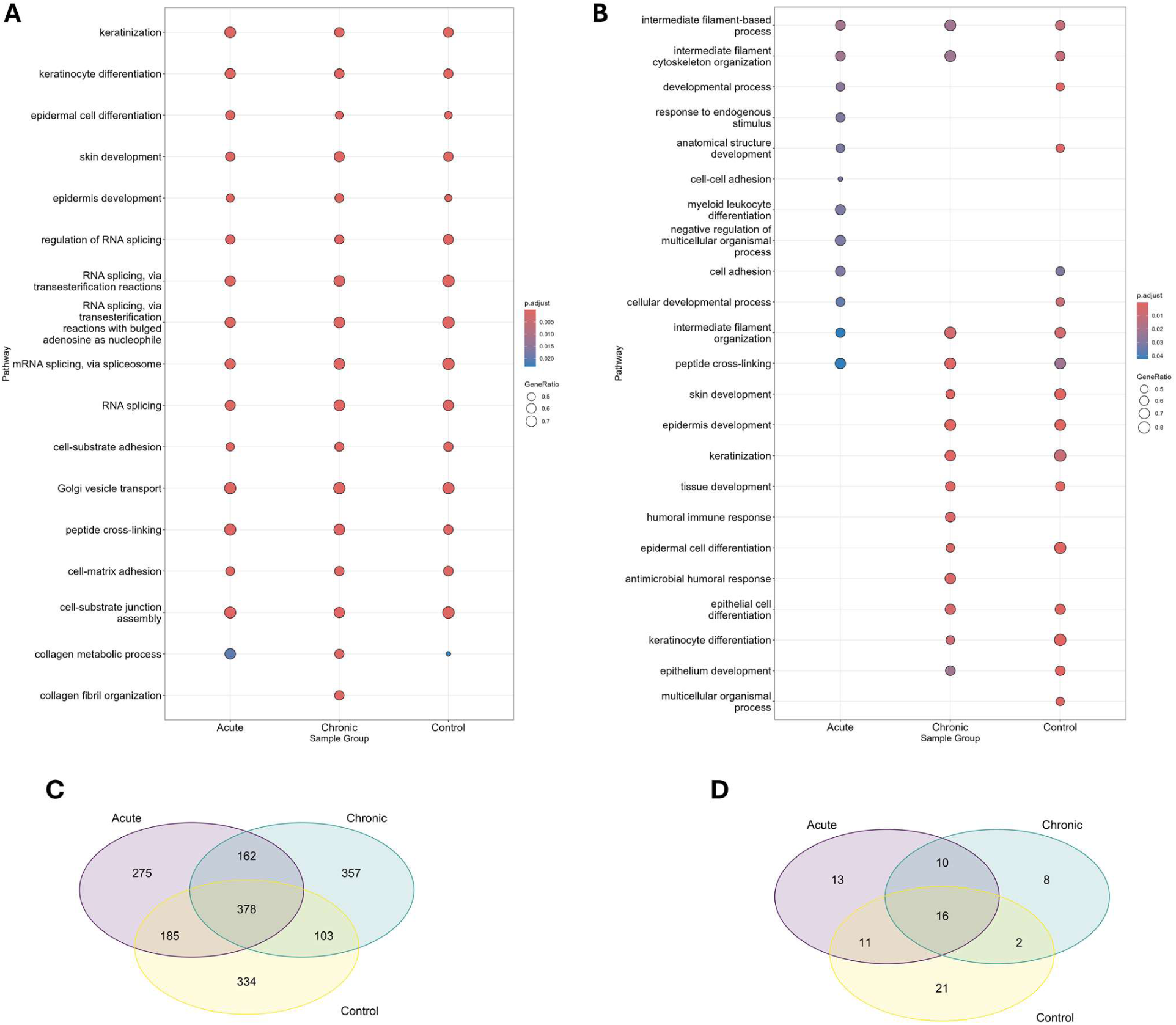
Gene Set Enrichment Analysis (GSEA) of the ranked row sum distribution of genes in Acute and Chronically wounded (n=3/group) and Control (n=2) skin samples. **A)** GSEA of all genes in the transcriptome per pathology group ranked on row sum distribution of the hypergraph, **B)** GSEA of top 1000 genes per pathology ranked on row sum distribution of the hypergraph, **C)** Venn diagram of the overlap of the top 1000 genes of the hypergraph row sum distribution per pathology group, D) Venn diagram of the overlap of the transcription factors mapped to the top 1000 genes of the hypergraph row sum distribution per pathology group.

Next, the top 1000 highly connected genes per sample group were compared to a list of known transcription factors. Fifty mapped to the acute samples, thirty-six to the chronic samples and fifty mapped to the control samples (Figure 4D). ORA analysis identified pathways in the acute samples such as integrated stress response signalling, maintenance of cell number and myeloid and mononuclear cell differentiation (all FDR <0.05), regulation of vasculature development, mononuclear cell differentiation and epithelial cell proliferation (all FDR <0.05) in the chronic samples and fat cell differentiation, response to leukaemia inhibitory factor and myeloid cell differentiation (all FDR <0.05) in control samples. Next, we analysed the set of transcription factors that were unique to each pathology. The acute samples expressed 13 unique transcription factors, including *HMG20B, USF2* and *TP53*. There were 8 transcription factors that were unique to the chronic wound samples. These included *ZEB2, MXD1* and *STAT2*. Finally, 21 genes were unique to the control samples, including *TFDP1, FOSL2*, and *JUN*. A list of transcription factors mapped to acute, chronic and control samples can be found in Supplementary Table S3.

#### Transcriptomic entropy analysis

There was a statistically significant difference in the overall transcriptomic entropy between the acute, chronic and control groups, with the control samples having the highest entropy, and the chronic samples having the lowest entropy (all p-adjusted <0.001) (Figure 5A).

**Figure 5:**
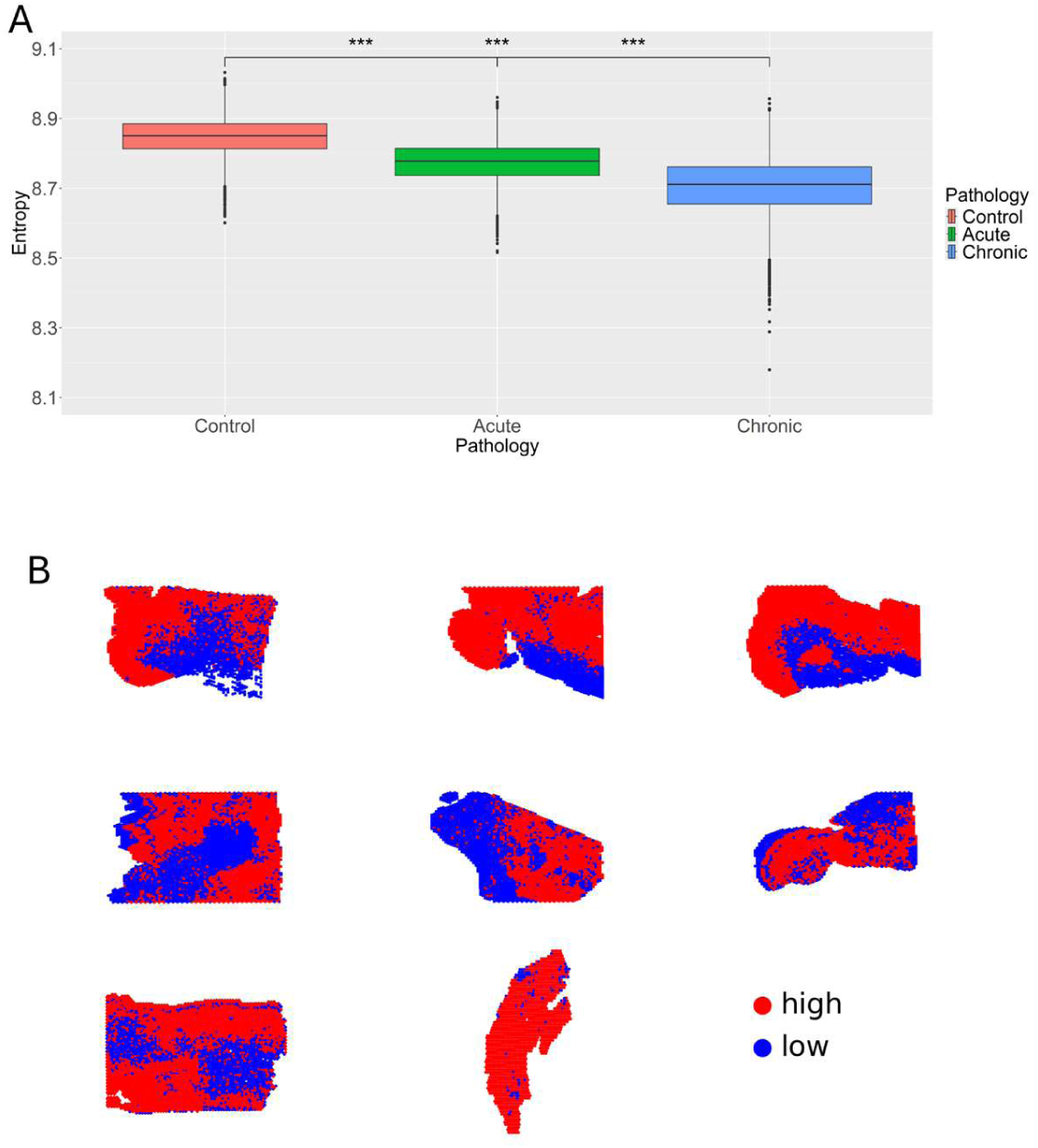
Transcriptomic entropy following hypergraph analysis of Acute (n=3) and Chronically (n=3) wounded samples and Control (n=2) skin samples. **A)** A statistically significant difference in the transcriptomic entropy calculated over the whole transcriptome per pathology group. Wilcoxon with Bonferroni correction, *** = P<0.0001. **B)** Entropy results categorised into high (≥9) and low (<9) entropy. All transcriptomic entropy calculated data subsetted for hypergraph analysis of a randomly selected 100 genes correlated to a randomly selected 1000 genes and iterated 10,000 times.

We next identified Leiden clusters of cell types which were shared between pathologies and compared their transcriptomic entropy. All comparisons between clusters containing keratinocytes between groups were statistically significant (p<0.001, Figure 6A). Additionally, all clusters containing fibroblasts, adipocytes and smooth muscle cells were also statistically different between groups (all p<0.001, Figure 6B&C). In addition, a number of the keratinocyte clusters in chronic wounds also showed low entropy across all three samples. When the transcriptomic entropy per cluster in each pathology group was split into “high” or “low” entropy, based on a cut off of 9, spatial differences became apparent. In all wounded samples, the wound bed and the layers directly underneath the wound were classed as having low entropy, whereas the remainder of the tissue had high entropy (Figure 5B). The median and p-values can be found in Supplementary Table S4. Boxplots displaying transcriptomic entropy of each Leiden cluster per pathology group can be found in Supplementary Figures 3, 4 and 5.

**Figure 6:**
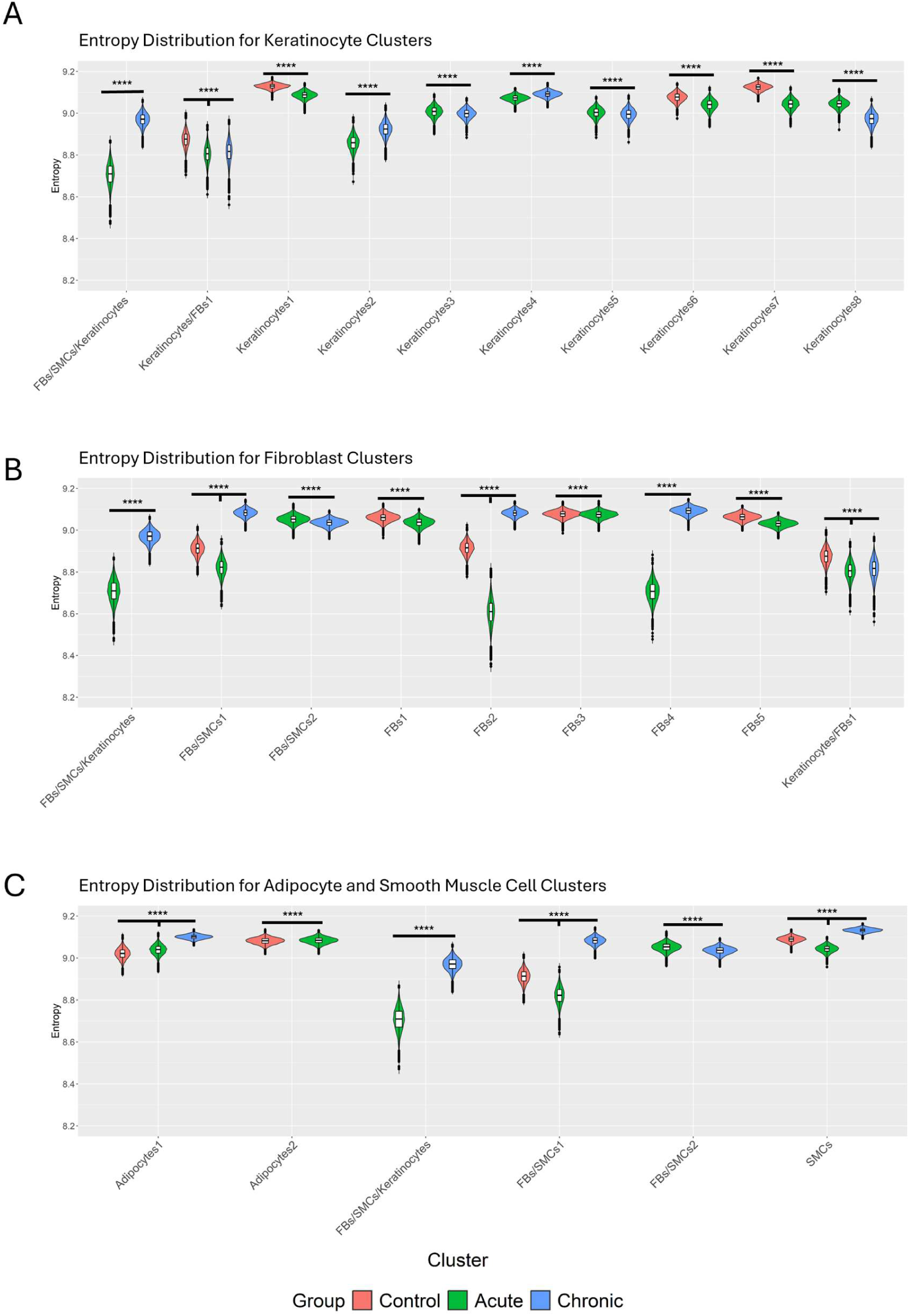
Transcriptomic entropy per Leiden cluster following hypergraph analysis of Acute (n=3) and Chronically (n=3) wounded samples and Control (n=2) skin samples. **A)** Entropy distribution in Leiden clusters containing keratinocytes. **B)** Entropy distribution in Leiden clusters containing fibroblasts. **C)** Entropy distribution in Leiden clusters containing adipocytes and smooth muscle cells. Wilcoxon performed on Leiden clusters between pathology groups, **** = p<2.2×10^−16^. A threshold was set of 90 spots per pathology group for a Leiden cluster to be considered present in the data. All transcriptomic entropy calculated data subsetted for hypergraph analysis of a randomly selected 100 genes correlated to a randomly selected 1000 genes and iterated 10,000 times. Leiden clusters had to be present in a minimum of two pathology groups to be included in the analyses. FBs: fibroblasts, SMC: smooth muscle cells

## Discussion

There have been very few studies that have evaluated the differences between human acute and chronic wounds mainly because the spectrum of disease is so diverse it would be challenging to encompass all wound phenotypes. The lack of tools that can provide meaningful comparisons across wound types has also been a limiting factor. However, tools available at present to specifically interrogate human tissue have provided new opportunities to evaluate chronic conditions hard to model in the laboratory. This study has used Visium Spatial Gene Expression to deconvolve the differences between acute and chronic wounds. All major cell types were identified within the data alongside marker genes used to distinguish between healed and non-healing wounds. The spatial relationship between Leiden clusters allowed for clusters relating to the wound bed to be identified, and spatial differences to be observed between the structure of the underlying cells in control and wounded skin samples. The analysis of higher order interactions through hypergraph modelling helped further deconvolve the key pathways associated with wound healing, whilst transcriptomic entropy analysis identified key differences between the order within the transcriptome that differentiated between acute and chronic wounds.

By creating one integrated dataset of all the samples, direct comparisons of Leiden clusters between healthy and wounded skin samples were possible. Twenty-nine Leiden clusters were identified, which, upon comparison with annotated H&E samples, broadly localised into the different cell layers expected in skin samples. To determine the cell types to be included in the clusters, the top differentially expressed genes were assessed using both The Human Protein Atlas and EnrichR. An advantage of using spatial gene expression is that the spatial location of the cell spots could also be taken into consideration. This improved the accuracy of the Leiden cluster annotations. However, Visium Spatial Gene Expression (V1) has a barcoded spot diameter of 55µM, with the centre of two neighbouring spots being approximately 100µM apart [30]. Whilst each capture area has 5000 barcoded spots, the size of the spots means that there is not the resolution to deconvolve single cells. A keratinocyte can have a diameter as small as 14µM [31], meaning multiple cells, of potentially different phenotypes and functions can be captured within a barcoded spot. Whilst this issue has now been mostly resolved with the new Visium HD, this version of the technology was not available when this study was conducted. To ensure that all cells present within a cell spot cluster were included, the transcriptomic profile of the Leiden cluster was investigated in enough detail to include multiple cell types in the assigning of its name.

When identifying the top 15 markers genes that differentiated between pathology groups, all were found in the dermis of the samples as opposed to the wound bed. This confirms previous knowledge that changes occurring deeper in the tissue are different between the chronic and acute samples, such as degradation of the ECM, unresolved inflammation and impaired angiogenesis leading to a hypoxic environment, further perpetuating the chronicity of wounds [32]. The changes known to occur in the ECM also support the genes identified as marking chronic wounds, including *COL1A1, FN1, BGN* and *POSTN*, alongside immune response genes *IGKC, IGHG* and *KRT16*. In contrast, the genes linked to immune response marking acute wound samples were focussed on the more acute inflammatory processes, such as complement activation (*C1QC* and *CFD*) which occurs shortly after injury [33]. However, persistent complement activation impairs wound healing and is therefore a target for inhibitors as a mechanism of treatment for chronic wounds [34]. The remainder of the genes marking acute samples were linked to cellular response to stress (*DUSP1*) and cell mobility and survival (*HSPA5* and *MARCK5*), both of which would be expected following injury, as immune cells, such as macrophages move into the wound bed to aid healing, and keratinocytes and fibroblast migrate into the area during the proliferative phase [32].

One aspect of wound healing that this study is unable to address is the effect of a patient’s treatment prior to the wound biopsy. Whilst samples were carefully selected based on their presentation of wounds, there are differences in the treatments the patients received. Whilst some received Negative Pressure Wound Therapy (for between 1 and 4 weeks), other wounds were only treated with Non-Antimicrobial Dressings. As this study was not designed to deconvolve the differences between dressing types, we are unable to discuss whether the dressing type has an impact on the results.

The use of spatial transcriptomic analysis on acute and chronic wound samples allows for description of phenotypically different keratinocyte and fibroblasts clusters and their spatial locations within the tissue to be identified. This information is lacking when scRNA sequencing data is used, although, Visium V1 does not have the resolution to identify individual cells. In the control samples, we were able to identify a normal structure of skin keratinocyte layers, with two keratinocyte clusters (7 and 8) forming the epidermis, Keratinocytes4 forming the basal epidermis, and the Keratinocytes10 being subepithelial. There were key differences between keratinocyte clusters seen within the wounds bed. In acute wound beds, Keratinocytes5, followed behind by Keratinocytes3 were present, with both having a marked decrease in the chronic samples but replaced with Keratinocytes2. The Keratinocytes2 cluster is observed in the middle layer of the acute samples, indicating either a movement of the cell types up towards the wound bed of the chronic samples, or the movement of the cells deeper into the tissue in acute samples. The most likely explanation is the upwards movement towards the skin surface, as part of the normal wound healing process of proliferation to replace cells lost during the injury [35]. Keratinocytes1 cluster was present in both the acute and control samples, but were almost entirely absent from the chronic samples, demonstrating some maintenance of normal tissue in acutely wounded samples.

There was much more variation seen in the clusters present in the middle layer of the samples. Whilst in control samples, there was a combination of fibroblast, keratinocytes and adipocyte clusters, there were only three fibroblast clusters, and an adipocyte cluster observed in the acute samples. All three adipocyte clusters (Adipocytes1-3), found in the middle and lower layers of acute and control samples, were almost entirely absent from the chronic wound samples. This finding is consistent with observations that there is a loss of fat underneath chronic wounds, and one arm of research is investigating the utilisation of adipose-derived stem cells to initiate cutaneous regeneration [36]. The presence of increased fibroblast clusters in chronic wound samples, would indicate that the fibrotic tissue is present, as would be expected in a non-healing wound [10]. This descriptive analysis allows for clear identification of clusters that are spatially distinct in acute, chronic and control samples. This work could be taken forward to aid either biomarker discovery, or for development of new therapeutics for chronic wounds.

The resolution of Visium V1 did not allow for identification of many immune cell specific clusters, however, by using marker genes typical of both pro- and anti-inflammatory processes, the differences between the acute and chronic samples were able to be assessed. Notable differences between the acute and chronic wounds were the presence of pro-inflammatory markers in both groups but with a difference in the expression of anti-inflammatory markers. In the acute wound bed, the expression was sparse, but the cell spots present were highly expressing anti-inflammatory markers, compared to the chronic wounds where there was more generalised expression. The inflammatory stage switches into the anti-inflammatory stage of wound healing at around 3 days post injury, with remodelling starting around 3 weeks post injury [37]. Our samples are between 8- and 10-days post injury, suggesting that there are very specific areas of anti-inflammatory activity within the wound bed in the stages following inflammation. The presence of anti-inflammatory markers within the chronic wound bed may be in agreement with the literature that it is an imbalance between pro- and anti-inflammatory that contributes to the chronicity of wounds [38]. The observed increase in proliferation in acute wound samples could relate to the presence of dendritic cells releasing growth factors to increase proliferation of keratinocytes which is observed within the literature [9].

We applied hypergraph analyses to measure higher order interactions [39], between genes to assess the organisation of the transcriptome, or between cell spot clusters in acute and chronic wound samples to understand spatial coordination of the tissue. Network structural analyses of the entire transcriptome revealed similarities between pathological groups per gene ontology analysis. This is likely due to the cellular functions, such as keratinisation and epidermis development occurring in all samples. However, when the data was filtered to the top 1000 highly ranked genes by row sum distribution, a measure of higher order interactions between genes, differences between the pathology groups were elucidated. This included pathways common to acute and control samples, which were missing in chronic samples, such as cell adhesion and cell developmental processes, and pathways common between chronic and control samples, such as epithelium development, and keratinisation. Despite this, pathways common between all three groups included intermediate filament cytoskeleton organisation, intermediate filament organisation and peptide cross-linking. This suggests that normal biological processes relating to skin homeostasis continue, even in the presence of an injury, regardless of whether the wound is acute or chronic.

To further deconvolve the genes with a high row sum and therefore high connectivity, which are likely to be functionally relevant, we mapped known transcription factors to the top 1000 genes by this metric. The transcription factors that uniquely mapped to the genes in each pathological group highlights biological pathways activated in each group. In the acute samples, *HMG20B* (required for complete cell division during mitosis [40]), *USF2* (a repressor for genes related to autophagy and lysosomes [41]), and *TP53* (regulation of cell homeostasis [42]) were identified. This compares to the unique chronic wound mapping transcription factors, including *ZEB2* (regulator of both epithelial-mesenchymal transition and mesenchymal-epithelial transition [43], *MXD1* (involved in the cMYC pathways of cellular proliferation and differentiation [44]) and *STAT2* (regulation of pro- and anti-inflammatory cellular responses [45]). These data show that fundamentally different pathways are important within the transcriptome of wounded skin, which could not be identified using traditional pathway analyses.

Entropy, a measure of disorder within the transcriptome, (alternatively can be thought of as the level of information content) [19], quantifies the network structure such that a cell spot with higher entropy, has greater heterogeneity in the higher order network structure. Transcriptomic entropy has been demonstrated to quantify functional organisational features relating to differentiation state [46]. The chronic samples had the lowest entropy overall, suggesting that the process of chronic disease fixes cells in a fully differentiated cell state, possibly contributing the pathogenesis of disease, the inability of the wound to heal and the formation of scar tissue which has yet to be remodelling [47]. The lower entropy observed in chronic wounds may also be due to the failure or inappropriate remodelling of the basement membrane beneath the wound bed, which usually occurs in healing wounds but is also disrupted in aging skin [48]. Interestingly, the keratinocyte clusters in the chronic wounds also showed low entropy, suggesting a distinct distraction from normal or acute wound healing behaviour. The control samples had the highest entropy overall, possibly linked to changing transcriptomic organisation in the normal cellular turnover found in healthy tissue. These findings suggest a gradation of cell entropy within wounded skin samples, where acute wounds maintain some of the cellular function of healthy skin, possibly linked to the normal structure of the skin layers underneath the wound bed, whereas chronic skin contains more cells that are fully differentiated or in a profibrotic state that they cannot return from [9]. Further analysis looking at the cluster level entropy per pathology identified more variation in the acute samples than both the chronic and control samples. This variation is likely to be indicative of the differences in cellular composition of the samples, and the different stages of wound healing present in the samples. Transcriptomic entropy analysis is rarely performed in pathological samples. The findings of a decrease in entropy in wounded samples compared to controls contrasts with entropy studies in cancer. The application of entropy analysis to published datasets with DNA copy number and a gene expression profile, revealed an increase in entropy in cancer samples compared to controls [49]. This difference between cancer and wound healing indicates that the changes that cause mutations in cellular DNA fundamentally change the transcriptomic entropy and increases the disorder or information content within a cell. These changes are not seen in samples undergoing wound healing.

This study used Visium Spatial Gene Expression to analyse acute and chronic wounds and compare them to healthy, unwounded skin samples. We used hypergraph analyses to provide new insights into the higher order coordination in wounded skin, and to identify genes that are highly coordinated within the transcriptome and the biological pathways that they are associated with. By measuring transcriptomic entropy, we were able to demonstrate increasing levels of order within the transcriptome from acute to chronic wound samples, with further deconvolution of the differences between acute and chronic wound possible when cell cluster-level transcriptomic entropy was assessed. By overlaying transcriptomic entropy onto the spatial images, identification of low entropy in and around the wound beds was possible. These analyses will provide a framework on which future biomarkers of chronic wounds can be identified. This will open the door to novel treatments targeted towards chronic wounds.

## Acknowledgments

We would like to thank Dr Leena Joseph for confirming our analysis of the pathology and structure of the samples.

## Author contributions

MCS: helped generate the initial transcriptomic samples, performed the analyses and wrote the manuscript

JAH: lead the generation of the initial transcriptomic data and chose the samples to be included in the study.

RM: contributed to figure generation, data analysis and proofread the manuscript.

LIW: contributed to figure generation, data analysis and proofread the manuscript.

TG: contributed to entropy and hypergraph analyses and contextualising the results.

SMB: provided advice on the analyses of the data.

AR: provided access to the samples from the Complex Wound@Manchester Biobank.

AS: conceptualised the project, provided supervision of the computational analysis and proofread the manuscript.

JW: conceptualised the project, provided clinical context to the data, provided clinical information on the samples and helped write the manuscript.

