## Supplemental Tables for "Spatial transcriptomic profiling reveals distinct signatures in acute versus chronic wounds through hypergraph modelling and transcriptomic entropy analysis"

**hapTable S1: Demographics of the Acute, Chronic and Control samples included in the study**

| **Sample** | **Stage** | **Sex** | **Ethnicity** | **Type of wound** | **Location** | **Wound age** | **Healed**  **(yes/no)** | **Comorbidity** | **Dressing** |
| --- | --- | --- | --- | --- | --- | --- | --- | --- | --- |
| Acute Disease 1 | Acute | M | White British | Traumatic wound sustained following fall from height, fasciotomy wound, sampling before grafting | Forearm | 8 days | Yes | None | Non-antimicrobial dressing |
| Acute Disease 2 | Acute | F | White British | Surgical wound, old traumatic wound debrided 10 days prior to sampling and reconstruction | Lower back | 10 days | Yes | Obesity | Negative pressure wound therapy for 2 weeks |
| Acute Disease 3 | Acute | M | Asian | Surgical wound, following flap reconstruction, debrided 1 week before sampling and grafting | Buttock | 7 days | Yes | G6P deficiency | Non-antimicrobial dressing |
| Chronic Disease 1 | Chronic | M | White British | Grade IV pressure ulcer secondary to poor mobility due to MS | Buttock | 18 months | No | Multiple sclerosis | Negative pressure wound therapy for less than 1 week |
| Chronic Disease 2 | Chronic | F | White British | Non-healing diabetic foot ulcer | Foot | 8 weeks | No | Hypertension, myocardial infarction, (previous bypass stent), non-insulin dependent diabetes, peripheral vascular disease, forefoot amputation | Non-antimicrobial dressing |
| Chronic Disease  3 | Chronic | F | White British | Lower limb traumatic wound, ORIF infected chronic non-healing ulcer | Ankle | 48 months | No | Hypertension | Non-antimicrobial dressing |
| Control 1 | Control | F | White British | Post-debridement tissue from unwounded area | Lower back | N/A | N/A | Obesity | Negative pressure wound therapy for 2 weeks |
| Control 2 | Control | F | White British | Fasciotomy wound from acute limb ischaemia secondary to occluded femoral popliteal bypass graft | Leg | N/A | N/A | Arterial vascular disease | Negative pressure wound therapy for 4 weeks |

**Table S2: Genes used to define anti- and pro-inflammatory states and immune cells.** Genes taken from a literature search.

| **State/ immune cell** | **Genes** |
| --- | --- |
| Pro-inflammatory_markers | IL1A, IL1B, CXCL8, IL6, IL10, TNF, IFNG, CCL2, HMGB1, HSPA1A, S100A8, IL13, CCL4 |
| Anti-inflammatory_markers | TGFB1, TGFB2, TGFB3, IL3, IL10, IL13, IL1RN, IL4 |
| Pro-inflammatory_macrophages | CD80, CD86, FCGR1A, FCGR2A, FCGR3A |
| Anti-inflamatory_macrophages | CD163, MRC1, ARG1, CLEC7A |
| General_macrophages | CD68, ITGAM |
| B-cells | PTPRC, CD19, MS4A1, CD79A, CD24, CD22 |
| T-activated | CD3D PTPRC, IL2RA |
| T-cytotoxic | CD8A, PTPRC, CD4 |
| T-helper | CD3D, PTPRC, CD4 |
| T-regulatory | CD3D, PTPRC, FOXP3 |
| Langerhans | CD207, CD1A |
| Dendritic_cells_type_1 | ITGAE, ITGAX, LY75, THBD, CD8A, BTLA, CADM1, CLEC9A |
| Dendritic_cells_type_2 | ITGAM, CD163, CD14, CD1C, SIRPA, CLEC10A, NOTCH2 |
| Proliferation | CCNB1, FOXM1, TOP2A, PCNA, MYBL2, BUB1, MKI67, PLK1, CCNE1, CCND1 |
| Angiogenesis | VEGFC, LYVE1, PECAM1, VEGFA, ANGPT1, ANGPT2, ENG, VCAM1 |

**Table S3: List of transcription factors in top 1000 highly connected genes in the whole transcriptome per pathology hypergraph.**

| **Pathology Group** | **Transcription factors** |
| --- | --- |
| Acute transcription factors | FOS, MYC, GATA3, KLF5, JUNB, CEBPA, TP63, RXRA, NFIX, KLF6, BCL11A, ZBTB7B, NFIB, EHF, CTBP1, SFPQ, HMG20B, GTF2I, EGR1, USF2, STAT6, TP53, STAT3, PHB2, RARG, RERE, NFAT5, KLF13, ELF1, RORA, HDGF, CEBPD, KLF4, HES2, AHDC1, TFCP2L1, MAZ, NR4A1, ZBTB4, ZBTB7A, ID3, NFE2L2, KLF10, BCL6, NFE2L1, FOSB, SKI, TRIM25, BCLAF1. |
| Acute unique transcription factors | HMG20B, USF2, TP53, RERE, KLF13, ELF1, AHDC1, MAZ, NR4A1, ZBTB4, NFE2L1, SKI, TRIM25. |
| Chronic transcription factors | TP63, MYC, KLF5, JUNB, KLF4, RUNX1, NFIB, CEBPA, HES2, TFCP2L1, EHF, NR2F2, ZEB2, ZBTB7B, SFPQ, FOS, GATA3, STAT3, NFIX, MXD1, NFE2L2, BHLHE40, RORA, CREB3L1, STAT6, ID3, XBP1, ETS1, CTBP1, CEBPD, ILK, STAT2, PHB2, GTF2I, EGR1. |
| Chronic unique transcription factors | NR2F2, ZEB2, MXD1, CREB3L1, XBP1, ETS1, ILK, STAT2. |
| Control transcription factors | FOS, KLF5, KLF4, FOSB, EGR1, JUNB, MYC, TP63, GATA3, RORA, JUN, RXRA, ZBTB7B, BCL11A, KLF6, BCL6, TFCP2L1, KLF10, MXI1, EHF, RARG, NFE2L2, ATF3, GTF2I, FOSL2, XRCC5, SNAI2, ZBTB7A, SMARCC1, MEF2A, HDGF, RUNX1, TFDP1, ATF4, SREBF2, PHB2, SP3, DR1, TAF7, POLR2A, ZKSCAN1, CBFB, NFIC, CEBPG, BHLHE40, TBL1XR1, NFIX, BCLAF1, NFAT5, KDM2A. |
| Control unique transcription factors | JUN, MXI1, ATF3, FOSL2, XRCC5, SNAI2, SMARCC1, MEF2A, TFDP1, ATF4, SREBF2, SP3, DR1, TAF7, POLR2A, ZKSCAN1, CBFB, NFIC, CEBPG, TBL1XR1, KDM2A. |

**Table S4: Median entropy values for acute, chronic and control entropy per cluster and comparison between pathological groups.** Wilcoxon test performed between acute and chronic, control and acute, and control and chronic samples, **** p< 2.2 x 10^−16^

| **Cluster** | **Acute Median Entropy** | **Chronic Median Entropy** | **Control Median Entropy** | **Acute_vs_Chronic** | **Control_vs_Acute** | **Control_vs_Chronic** |
| --- | --- | --- | --- | --- | --- | --- |
| **Adipocytes1** | 9.04 | 9.1 | 9.02 | **** | **** | **** |
| **Adipocytes2** | 9.08 | NA | 9.08 | NA | **** | NA |
| **FBs/SMCs/Keratinocytes** | 8.71 | 8.97 | NA | **** | NA | NA |
| **FBs/SMCs1** | 8.82 | 9.08 | 8.91 | **** | **** | **** |
| **FBs/SMCs2** | 9.05 | 9.04 | NA | **** | NA | NA |
| **FBs1** | 9.04 | NA | 9.06 | NA | **** | NA |
| **FBs2** | 8.61 | 9.08 | 8.92 | **** | **** | **** |
| **FBs3** | 9.08 | NA | 9.08 | NA | **** | NA |
| **FBs4** | 8.71 | 9.09 | NA | **** | NA | NA |
| **FBs5** | 9.03 | NA | 9.06 | NA | **** | NA |
| **Keratinocytes/FBs1** | 8.81 | 8.82 | 8.88 | **** | **** | **** |
| **Keratinocytes1** | 9.09 | NA | 9.13 | NA | **** | NA |
| **Keratinocytes2** | 8.86 | 8.92 | NA | **** | NA | NA |
| **Keratinocytes3** | 9.01 | 9.0 | NA | **** | NA | NA |
| **Keratinocytes4** | 9.07 | 9.09 | NA | **** | NA | NA |
| **Keratinocytes5** | 9.0 | 9.0 | NA | **** | NA | NA |
| **Keratinocytes6** | 9.04 | NA | 9.08 | NA | **** | NA |
| **Keratinocytes7** | 9.04 | NA | 9.13 | NA | **** | NA |
| **Keratinocytes8** | 9.05 | 8.97 | NA | **** | NA | NA |
| **SMCs** | 9.04 | 9.13 | 9.09 | **** | **** | **** |
