## Supplemental Figures for "Spatial transcriptomic profiling reveals distinct signatures in acute versus chronic wounds through hypergraph modelling and transcriptomic entropy analysis"

### Slide 1
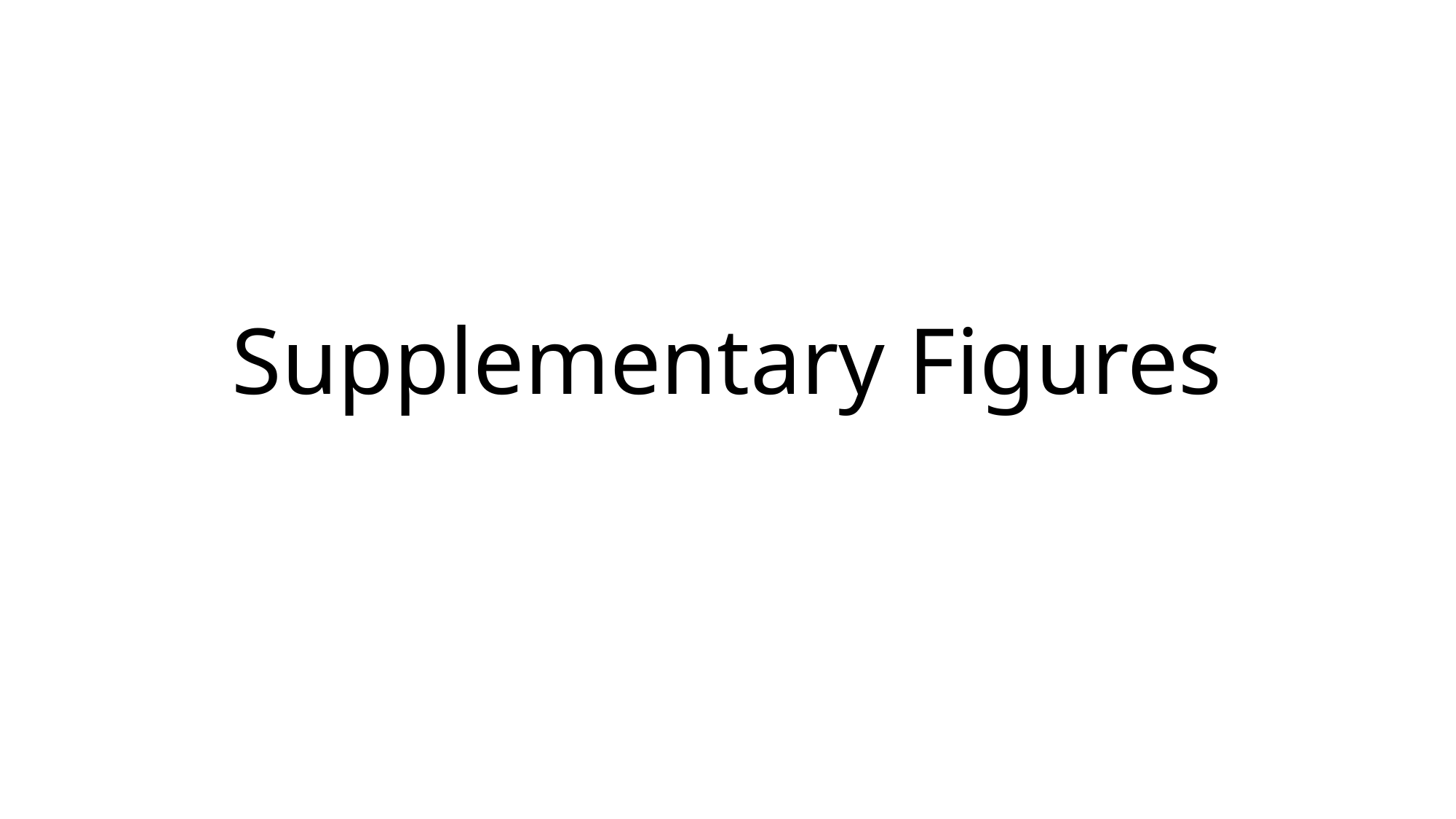

#### Supplementary Figures

### Slide 2
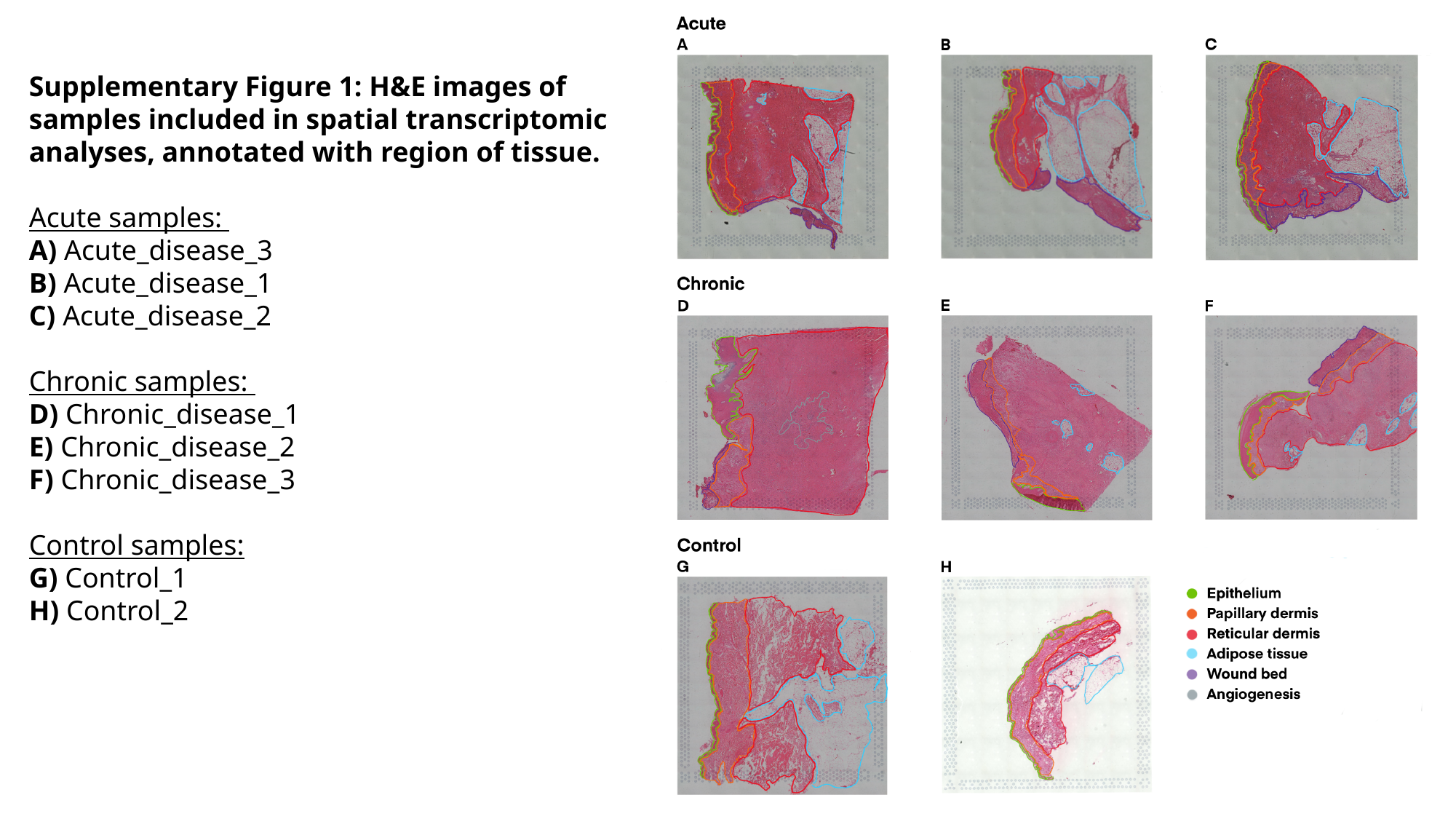

Supplementary Figure 1: H&E images of samples included in spatial transcriptomic analyses, annotated with region of tissue.
Acute samples:
A) Acute_disease_3
B) Acute_disease_1
C) Acute_disease_2
Chronic samples:
D) Chronic_disease_1
E) Chronic_disease_2
F) Chronic_disease_3
Control samples:
G) Control_1
H) Control_2

### Slide 3
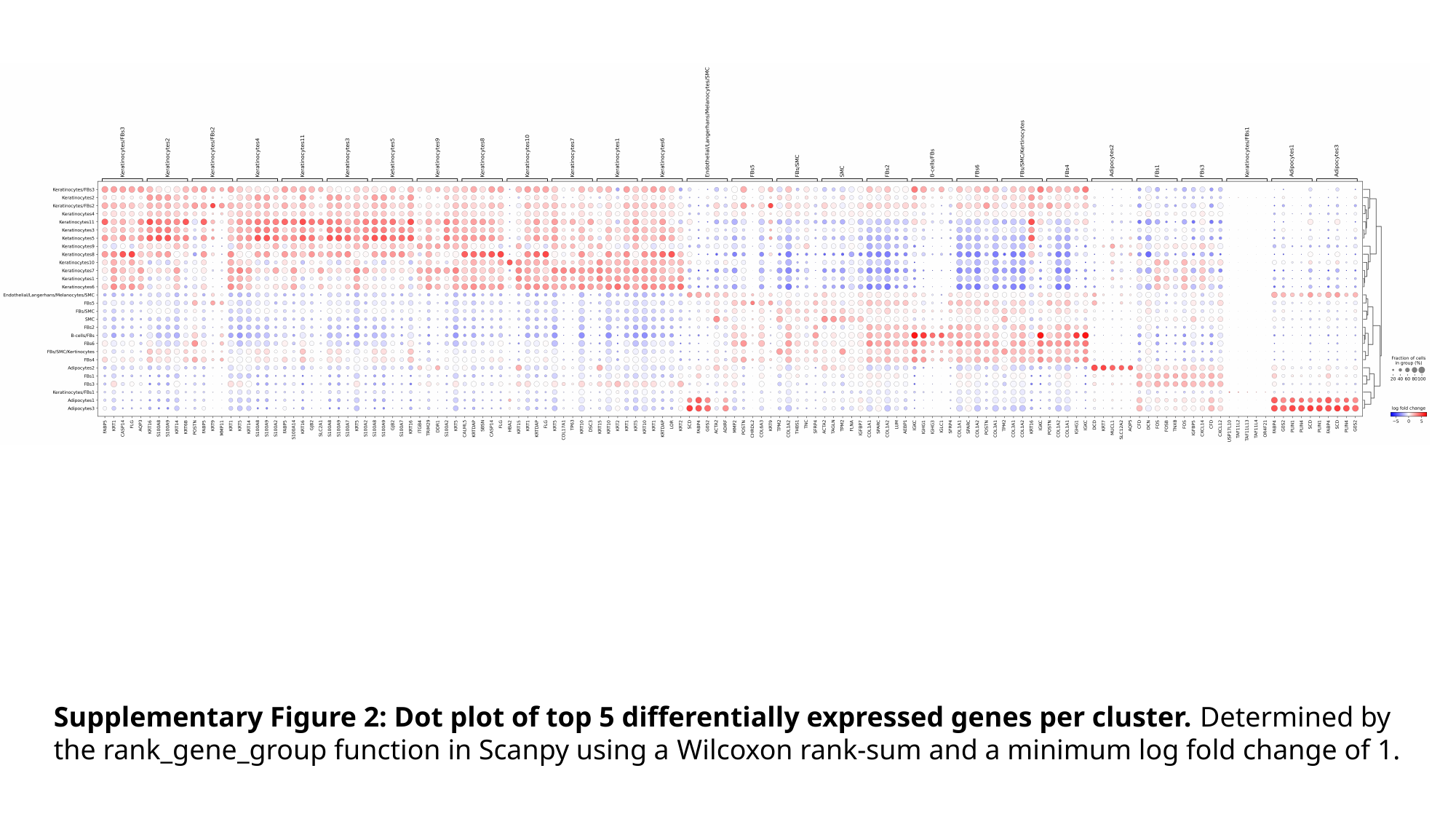

Supplementary Figure 2: Dot plot of top 5 differentially expressed genes per cluster. Determined by the rank_gene_group function in Scanpy using a Wilcoxon rank-sum and a minimum log fold change of 1.

### Slide 4
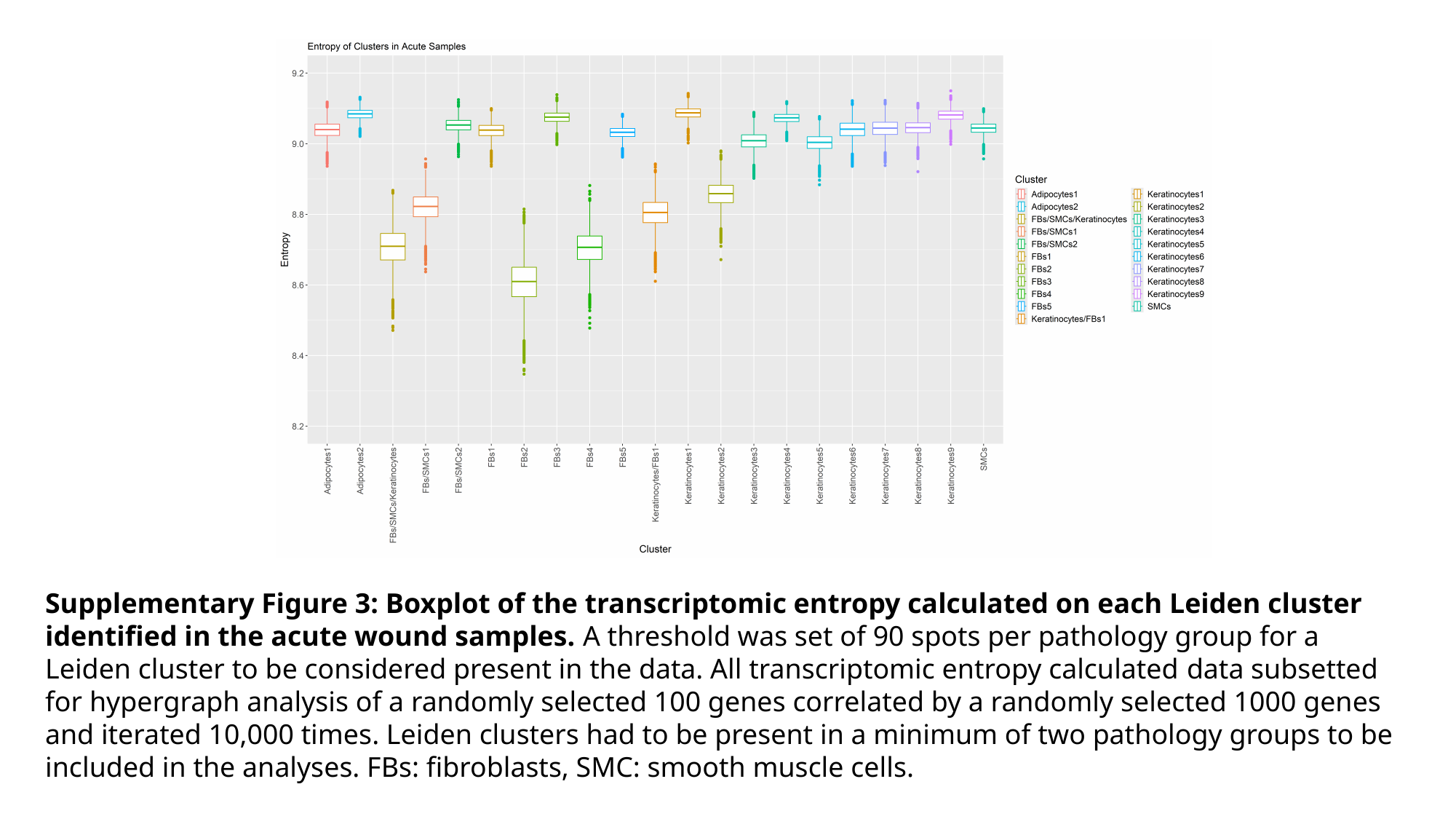

Supplementary Figure 3: Boxplot of the transcriptomic entropy calculated on each Leiden cluster identified in the acute wound samples. A threshold was set of 90 spots per pathology group for a Leiden cluster to be considered present in the data. All transcriptomic entropy calculated data subsetted for hypergraph analysis of a randomly selected 100 genes correlated by a randomly selected 1000 genes and iterated 10,000 times. Leiden clusters had to be present in a minimum of two pathology groups to be included in the analyses. FBs: fibroblasts, SMC: smooth muscle cells.

### Slide 5
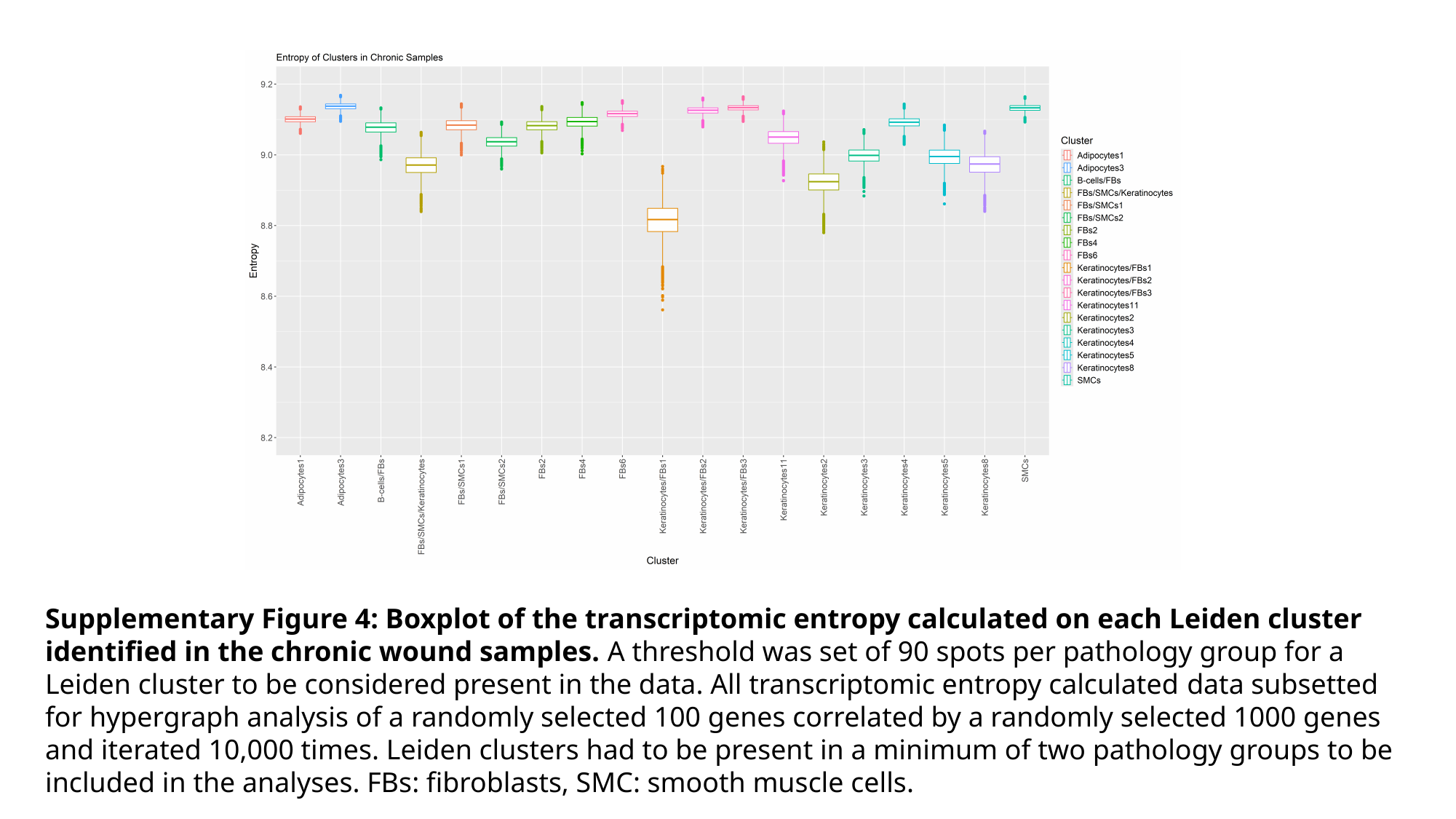

Supplementary Figure 4: Boxplot of the transcriptomic entropy calculated on each Leiden cluster identified in the chronic wound samples. A threshold was set of 90 spots per pathology group for a Leiden cluster to be considered present in the data. All transcriptomic entropy calculated data subsetted for hypergraph analysis of a randomly selected 100 genes correlated by a randomly selected 1000 genes and iterated 10,000 times. Leiden clusters had to be present in a minimum of two pathology groups to be included in the analyses. FBs: fibroblasts, SMC: smooth muscle cells.

### Slide 6
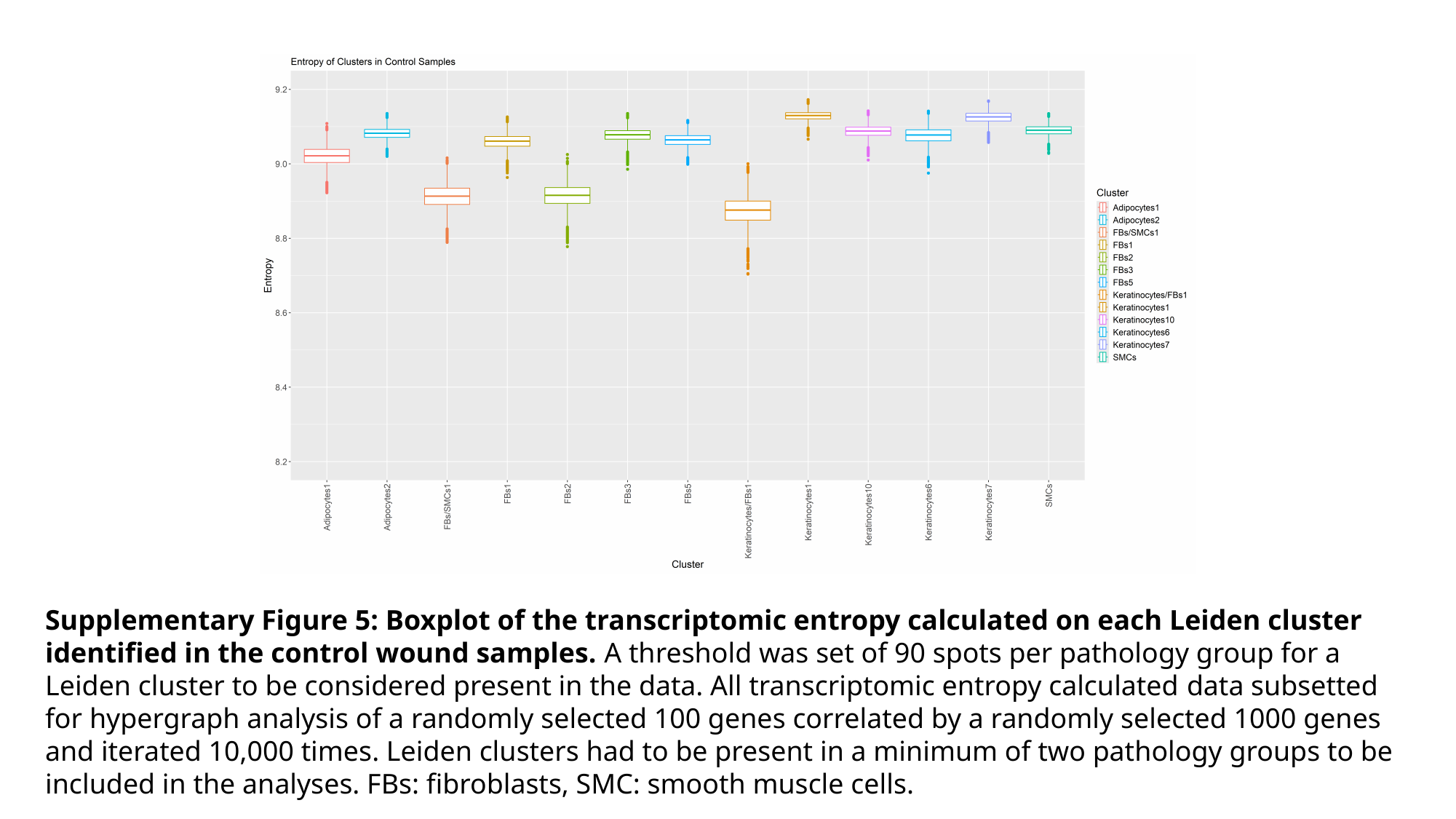

Supplementary Figure 5: Boxplot of the transcriptomic entropy calculated on each Leiden cluster identified in the control wound samples. A threshold was set of 90 spots per pathology group for a Leiden cluster to be considered present in the data. All transcriptomic entropy calculated data subsetted for hypergraph analysis of a randomly selected 100 genes correlated by a randomly selected 1000 genes and iterated 10,000 times. Leiden clusters had to be present in a minimum of two pathology groups to be included in the analyses. FBs: fibroblasts, SMC: smooth muscle cells.
